# No evidence for the presence and control of tumor growth by cytoplasmic stress granules in pancreatic cancer

**DOI:** 10.1101/2023.11.13.566518

**Authors:** Maxime Libert, Jean Fain, Christelle Bouchart, Tatjana Arsenijevic, Jean-Luc Van Laethem, Patrick Jacquemin

## Abstract

Cytoplasmic stress granules (SG) are attracting growing attention in cancer research^1^. As such, a study published in Cell^2^ has certainly served as a trigger for this trend which currently asserts that SG are a relevant therapeutic target. Here, we would like to express a warning about the conclusions of this article, as well as of the subsequent study^3^, and more generally on a methodological approach frequently used in research studying SG in cancer and which we believe to be misleading.

---

The study mentioned above^2^ claims that SG confers cytoprotection against stress stimuli and chemotherapeutic agents in mutant KRAS cancers. The results first indicate that SG are markedly elevated in arsenite-treated mutant KRAS cell lines compared to arsenite-treated wild-type KRAS cell lines (with a SG index 12 times higher in the former). With such a difference, mutant KRAS cell lines should have been spontaneously selected in the past by SG researchers for their studies; however, and even if a founder effect should be considered, this is not the case as 17 out of 21 cell lines subsequently used in top 10 articles from PubMed search with “stress granules cell lines” (sorted by Best Matches on 03/14/2023) are wild-type KRAS. To clarify this point, we subjected wild-type and mutant KRAS cell lines to arsenite treatment, in the same way as the previous study. SG were detected by G3BP1 and CAPRIN1 immunolabelling. The calculated SG index revealed no correlation between the SG index and the presence of KRAS mutations (Figure 1A).

**Figure 1.**
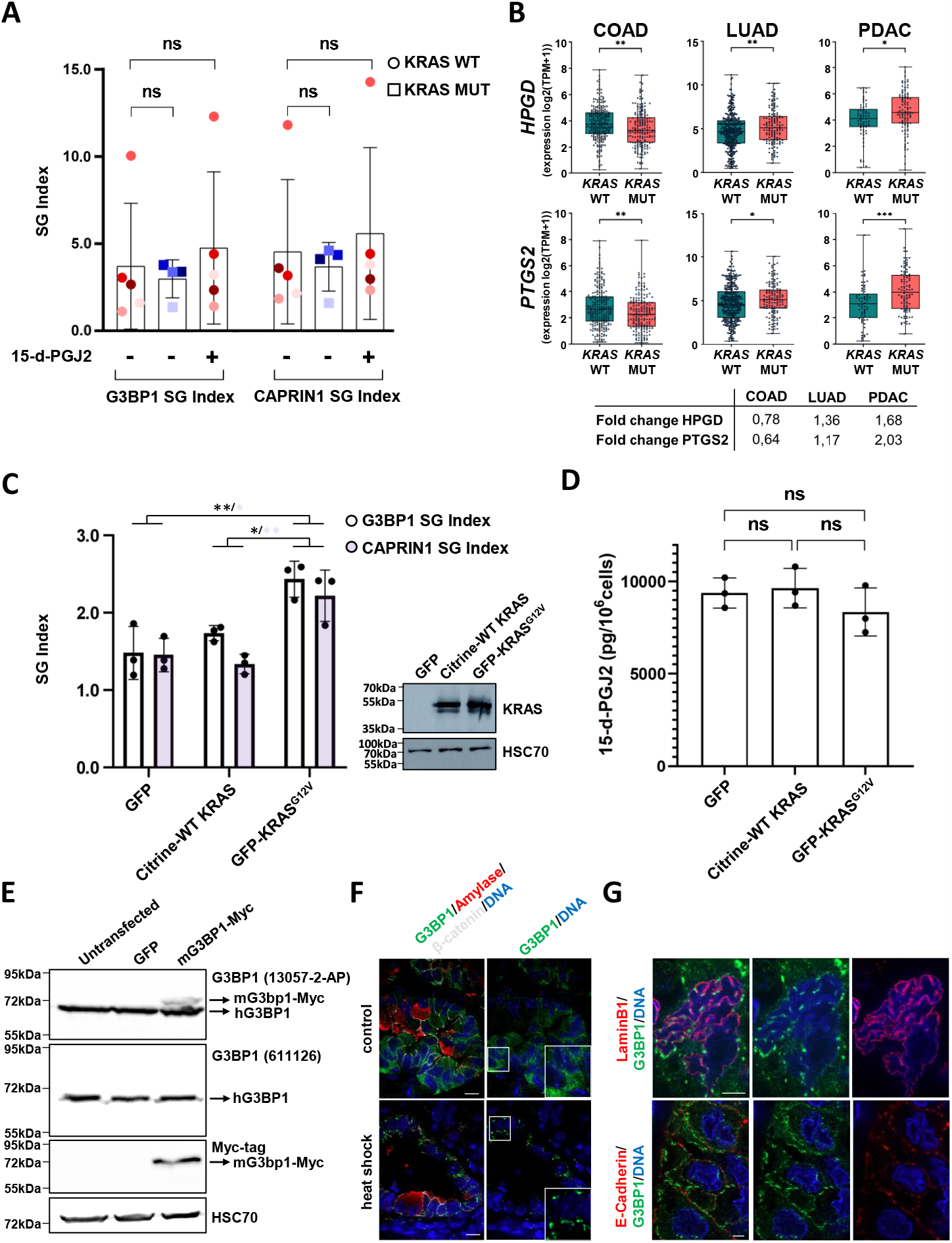
Kras mutation does not influence SG formation and cytoplasmic stress granules are not present in pancreatic cancer. (**A**) Five wild-type and four mutant KRAS cell lines were subjected to arsenite treatment to induce cytoplasmic stress granule (SG) formation. SG were detected by G3BP1 and CAPRIN1 immunolabelling and quantified by calculating the SG index (SG surface divided by cell surface). The data are presented as the percentage of SG surface relative to the total cell surface. (**B**) The upper panels represent mRNA expression levels of *HPGD* and *PTGS2* in three human cancers (colon adenocarcinoma-COAD, lung adenocarcinoma-LUAD, and pancreatic ductal adenocarcinoma-PDAC) categorized by Kras mutation status. Each dot represents one patient. The y-axis represents RNA-Seq by expression log_2_(TPM+1). The lower panel shows the fold changes between mean expressions in *KRAS* WT and *KRAS* mutant patients. (**C**) HEK293 cells were transfected with plasmids encoding GFP, Citrine-wild-type (WT) KRAS and GFP-KRAS^G12V^, and treated as described in A. SG index was calculated as described in A. To confirm transfection efficiency, whole cell lysates were collected and subjected to Western blotting with anti-KRAS antibody, HSC70 serving as a loading control. (**D**) Levels of 15-d-PGJ2 secretion were measured in HEK293 cells transfected as described in C. (**E**) Western blotting performed on protein extracts from untransfected, GFP-transfected, and mouse (m) G3BP1-transfected HeLa cells with G3BP1 (13057-2-AP), G3BP1 (611126), and Myc-Tag antibodies. Human (h) G3BP1 was detected by both G3BP1 antibodies, whereas mG3BP1 was only recognized by the former. (**F**) Sections from cerulein-treated ElaK pancreas shocked at 37°C (control) or 43°C (heat shock) during 30 minutes in PBS were immunolabelled with G3BP1, amylase (acinar cell marker) and β-catenin (plasma membrane marker). Nuclei were counterstained with Hoechst. Scale bar: 10 μm. (**G**) Sections from human PDAC were immunolabelled with G3BP1, LaminB1 (nuclear lamina marker) and E-cadherin (plasma membrane marker). Nuclei were counterstained with Hoechst. Scale bar: 5 μm. (**A, C, D**) Data presented are from a representative experiment out of at least 3 independent experiments leading to the same conclusion. The results are presented as means ± standard error of the mean (SEM). Quantifications were performed on 10 randomly acquired images at 100x magnification for each cell line. *, p < 0.05; **, p < 0.01; ***p <0.001; ns, not significant.

Another conclusion^2^ states that mutant KRAS upregulates SG by stimulating the production of prostaglandin 15-d-PGJ2. To reach this conclusion, it was shown that 15-d-PGJ2 obviates the requirement of mutant KRAS for the upregulation of SG in wild-type KRAS lines. To evaluate this result, we treated wild-type KRAS cells with 15-d-PGJ2, as previously described^2^. Again, we observed no difference in the SG index between untreated and prostaglandin-treated cells (Figure 1A). It also appeared *in vitro* that mutant KRAS is involved in the metabolism of 15-d-PGJ2 by increasing *PTGS2* mRNA (COX-2, the enzyme producing 15-d-PGJ2) and by decreasing *HPGD* mRNA (the catabolic enzyme), and that in humans, an enrichment of *PTGS2* was observed in lung adenocarcinoma (LUAD) with KRAS mutations compared to LUAD with wild-type KRAS^2^. However, a comparison including a much larger number of patients was unable to confirm that KRAS mutations control in opposite direction the expression of these two enzymes in LUAD, as well as in colon adenocarcinoma and pancreatic ductal adenocarcinoma, two other cancers with highly prevalent KRAS mutations (Figure 1B), casting doubt on whether this pathway is a regulator of SG in cancer. In addition, mutant KRAS has been shown to exert cell non-autonomous control on SG index through 15-d-PGJ2^2^. To verify this conclusion, we transfected wild-type KRAS cells with a GFP, wild-type, or mutant KRAS expression vector and measured 15-d-PGJ2 in the culture medium; though we observed a weak 1.5-fold increase in SG index in the presence of mutant KRAS (Figure 1C), far from the 5-fold increase previously described^2^, we were unable to detect any 15-d-PGJ2 increase in the culture medium of cells transfected with mutant KRAS, compared to that of cells transfected with GFP or wild-type KRAS (Figure 1D). Thus, our results do not confirm the observations that mutant KRAS stimulates 15-d-PGJ2 production, and that 15-d-PGJ2 upregulates SG.

Furthermore, we found that SG detection in mouse neoplastic lesions^2^ relies on the use of a monoclonal anti-G3BP1 antibody that recognizes the human form of G3BP1, but not the mouse form, possible because the immunogenic sequence (from human G3BP1) used to generate this antibody shows discrepancies in its mouse counterpart. We experimentally confirmed this information, both by Western blotting (Figure 1E) and immunocytolabeling (data not shown), which indicates that the previous detection of SG in mouse neoplasia^2^ is likely artefactual. Using a polyclonal antibody recognizing mouse G3BP1, we were unable to detect SG in mouse neoplasia whereas SG were detectable in mouse neoplasia subject to heat shock (Figure 1F).

Cytoplasmic SG have also been described in human pancreatic cancer specimens^2^. In addition, a subsequent study claims that cytoplasmic SG are upregulated in obesity-associated PDAC which is dependent on SG for its accelerated growth, compared to a PDAC not associated with obesity^3^. A key result of this study shows using G3BP1 immunolabelling that cytoplasmic SG are present in pancreatic cancer cells, and we strikingly noted that some were located in their nucleus^3^ (an organelle where cytoplasmic SG are absent). We sought to confirm these results, but came to a different conclusion. We found that G3BP1 was present diffusely in the cytoplasm of human PDAC cells, although in some areas G3BP1 condensates were also present (Figure 1G), and this independently of whether the PDAC is associated with obesity or not. In these areas, colabelling experiments showed that G3BP1 was mainly localized to the inner nuclear membrane and the plasma membrane of PDAC cells, cellular localizations that do not correspond to the presence of G3BP1 in cytoplasmic SG (Figure 1G). Again, our results do not support that cytoplasmic SG are present in pancreatic tumor cells and play a role in tumorigenesis.

In a more general way, studies focusing on SG in cancer are frequently based on experiments where cell lines are treated with arsenite to induce SG formation. This is because arsenite treatment generates oxidative stress like that which may occur in cancer cells. However, arsenite (at the concentration used to produce SG) rapidly arrests protein synthesis, raising questions on the relevance of the conclusions drawn from these experiments in the cancer field, as it is conceptually complicated to conceive that a biological mechanism identified *in vitro* in arsenite-treated cells can be preserved *in vivo* in cancer cells of obese patients or responding to chemotherapeutic treatment by producing SG (and consequently by no longer synthesizing proteins). Furthermore, to the best of our knowledge, the presence of stress granules has very rarely been described in tumors^4-5^, including a form of medulloblastoma with a mutation in a protein that promotes SG assembly^5^. In addition, there are no studies describing the presence of SG in samples from patients (or mouse models) treated with chemotherapy, regardless of the drug with reactive oxygen species-producing activity used. In this context, we conclude that SG do not promote tumorigenesis and cannot be considered a legitimate therapeutic target in cancer.

## Materials and methods

### Mouse model and treatments

All procedures described in this study were performed with the approval of the animal welfare committee of the UCLouvain Health Sciences Sector (Brussels, Belgium; ethic number: 2021/UCL/MD/054). ElastaseCre^ER^ LSLKras^G12D^ (ElaK) mice were maintained in a CD1-enriched background; this mouse model allows the expression of a mutated Kras^G12D^ allele specifically in acinar cells. 6-week-old ElaK mice were first treated with tamoxifen (T5648-1G, Sigma Aldrich. 30 mg/ml in corn oil) by oral gavage and 4-hydroxytamoxifen (H7904-25mg, Sigma Aldrich. 0.3 mg/ml in corn oil) by subcutaneous injection. Three sets of tamoxifen and 4-hydroxytamoxifen treatments, each separated by 48 hours, allow the recombination of the LSL cassette and the expression of Kras^G12D^ from its respective endogenous locus. To induce acute pancreatitis, 8-week-old tamoxified ElaK mice received 7 hourly intraperitoneal injections of cerulein (AS-24252, Eurogentec. 100-150 μl in phosphate buffered saline-PBS, pH 7.4, 125 μg/kg), every other day, for 5 days. Mice were sacrificed by cervical dislocation and pancreas were collected for subsequent analyses.

### Heat shock

For heat shock, mice were anesthetized and their pancreas was carefully displaced outside the abdominal cavity. The exposed pancreas was then incubated in PBS at 37°C or 43°C for 30 minutes. After sacrifice, the pancreas was fixed in 4% paraformaldehyde (HT501128, Sigma Aldrich) at 4°C for 4 hours, before embedding in paraffin.

### Cell lines

Cells were cultured in a humidified 5% CO_2_ incubator at 37°C in the conditions described below. KPE cells were derived from a tumour induced in ElaK p53^R172H^ mice.

| Cell line | Medium | Antibiotic | Supplement |
| --- | --- | --- | --- |
| HeLa | DMEM | 50 U/ml Pen. and 50 µg/ml Strep. | 10% FBS |
| SkChA1 | DMEM | 50 U/ml Pen. and 50 µg/ml Strep. | 10% FBS; 1mM Sodium pyruvate; 2 mM L-Glutamine |
| BXPC3 | RPMI | 50 U/ml Pen. and 50 µg/ml Strep. | 10% FBS |
| U2OS | DMEM | 50 U/ml Pen. and 50 µg/ml Strep. | 10% FBS |
| HEK293 | DMEM | 50 U/ml Pen. and 50 µg/ml Strep. | 10% FBS |
| EGI-1 | DMEM | 50 U/ml Pen. and 50 µg/ml Strep. | 10% FBS; MEM NEAA, 2 mM L-Glutamine |
| Panc1 | DMEM | 50 U/ml Pen. and 50 µg/ml Strep. | 10% FBS; 1mM Sodium pyruvate; 2 mM L-Glutamine |
| HPAC | DMEM | 50 U/ml Pen. and 50 µg/ml Strep. | 10% FBS |
| KPE | DMEM | 50 U/ml Pen. and 50 µg/ml Strep. | 10% FBS |

DMEM = Dulbecco's modified Eagle medium (61965026, Life Technologies)
RPMI = Roswell Park Memorial Institute (21875034, Life Technologies)

DMEM = Dulbecco’s modified Eagle medium (61965026, Life Technologies)

RPMI = Roswell Park Memorial Institute (21875034, Life Technologies)

MEM NEAA = Minimum Essential Medium Non-Essential Amino Acids (11140-035, Life Technologies)

FBS = Fetal bovine serum (F9665-500ML, Sigma Aldrich)

Pen. Strep. = Penicillin-Streptomycin (15070063, Life Technologies) Sodium Pyruvate (11360070, Life Technologies, Belgium)

L-Glutamine (25030081, Thermo Fisher Scientific)

### Stress treatments

Formation of SG was induced by incubating cells with 0.1 mM sodium arsenate for 30 minutes. Cells were treated with 50 μM 15-d-PGJ2 for 60 minutes.

### Transfection

30,000 cells were seeded in a 24-well plate for SG quantification or 10^6^ cells in 10-cm diameter dish for validation of transfection efficiency by Western blot. Calcium phosphate transfection^6^ was performed with 0.21 μg DNA/cm^2^. The plasmid DNA used for transfection was pEGFP, Citrine-KRAS^7^ and GFP-KRAS^G12V 8^. They were amplified in competent *Escherichia Coli* (L2005, Promega) following the manufacturer’s instructions. The plasmids were purified using Pure Yield™ Plasmid Midiprep system (A2495, Promega) and their concentration were determined using NanoDrop™ One (Thermo Fisher Scientific).

### SG quantification

SG quantifications were performed on 10 randomly acquired images at 100x magnification per replicate for each cell line. The Halo software (Indica Labs, v.3.3.2541) was employed for the quantification of stress granules. Using an algorithm (Indica Labs – Object colocalization FL v1.0), a threshold was set to distinguish stress granules from background noise, and the software automatically detected and quantified individual granules based on size, intensity, and shape parameters. Using another algorithm (Indica Labs – Area quantification FL v2.1.7), a second threshold was set to determine the cytoplasmic area of the cells. The stress granule index was calculated by dividing the total surface occupied by stress granules by the total surface of the cells in each image. This calculation was performed for every image, and the average value was obtained by combining the results from all images.

### Western blotting

10^6^ cells were seeded in 10-cm diameter dish. Cells were lysed by vortexing and repetitive pipetting in a buffer composed of 50 mM Tris-Cl, 150 mM sodium chloride, 0.25% sodium deoxycholate, 1% NP-40, 1 mM sodium orthovanadate, 2% sodium dodecyl sulfate (SDS), containing a protease inhibitor cocktail (11836153001, Sigma Aldrich). Lysates were maintained on ice during the procedure. Then, cell debris were pelleted by centrifugation (14,000 *g*, 10 minutes, 4°C). Proteins were quantified using a Bradford assay (23200, Thermo Fisher Scientific). Lysates containing 50 μg total proteins were separated on 12.5% SDS polyacrylamide gels. Polyvinylidene difluoride membranes (ISEQ00010, Millipore) were blocked with a solution of 5% low-fat milk diluted in Tris buffered saline (TBS)/0.1% Tween-20 (P2287, Sigma-Aldrich) for 1 hour at room temperature (RT). Membranes were incubated overnight at 4°C with primary antibodies against KRAS4B (WH0003845M1, Sigma Aldrich, 1 μg/ml) or HSC70 (sc-7298, Santa Cruz, 40 ng/ml) used as loading control, diluted in blocking buffer. Then, membranes were washed with TBS/0.1% Tween-20 and incubated with secondary antibodies for 1 hour at RT. After incubation, membranes were washed with TBS/0.1% Tween-20 and signals were revealed using SuperSignal™ West Pico PLUS Chemiluminescent Substrate (34577, Thermo Fisher). Pictures were taken with the Amersham ImageQuant 800 imaging system (Cytiva).

### Human pancreas specimens

The use of human tissue with histologically confirmed pancreatic ductal adenocarcinoma and without neoadjuvant treatment was approved by the Committee of Medical Ethics—Erasme Hospital (P2021/382). BMI data was obtained for each patient.

### Immunofluorescence

60,000 cells were seeded on a coated-polylysin (P2636, Sigma Aldrich - 50μg/ml in PBS, 25 min, 37°C) coverslip in 24-well plate. One day later, cells were fixed with 4% paraformaldehyde at 4°C. Six-μm paraffin-embedded tissue sections were deparaffinized and antigen retrieval was performed by heating the slides for 20 minutes in Tris-EDTA buffer (pH 9) using Lab Vision PT Module (Thermo Fisher Scientific). Sections and coverslips were rinsed 5 minutes in PBS, and permeabilized in PBS, 0.3% Triton X-100 (3051.3, Carl Roth) for 5 minutes at RT. After permeabilization, samples were blocked in PBS/3% low-fat milk/10% bovine serum albumin (A906-100G, Sigma Aldrich)/0.3% Triton X-100 (blocking buffer) for 45 minutes at RT. Primary antibodies were diluted in blocking buffer and incubated with the samples overnight at 4° C. Primary antibodies and antibody dilution are listed below. Then, samples were washed with 0.1% Triton X-100 in PBS. Secondary antibodies were diluted in PBS, 10% bovine serum albumin, 0.3% Triton X-100, applied at dilution listed below, and incubated for 1 hour at 37° C. Nuclei were counterstained with Hoechst (B2261, Sigma Aldrich – 3.2 μM). Then, samples were washed with 0.1% Triton X-100 in PBS and the slides were mounted in fluorescence mounting medium (S302380-2, Agilent). Imaging was performed with Cell Observer Spinning Disk Confocal Microscope using the ZEN software. Laser power and exposure times were similar for images from each data sets.

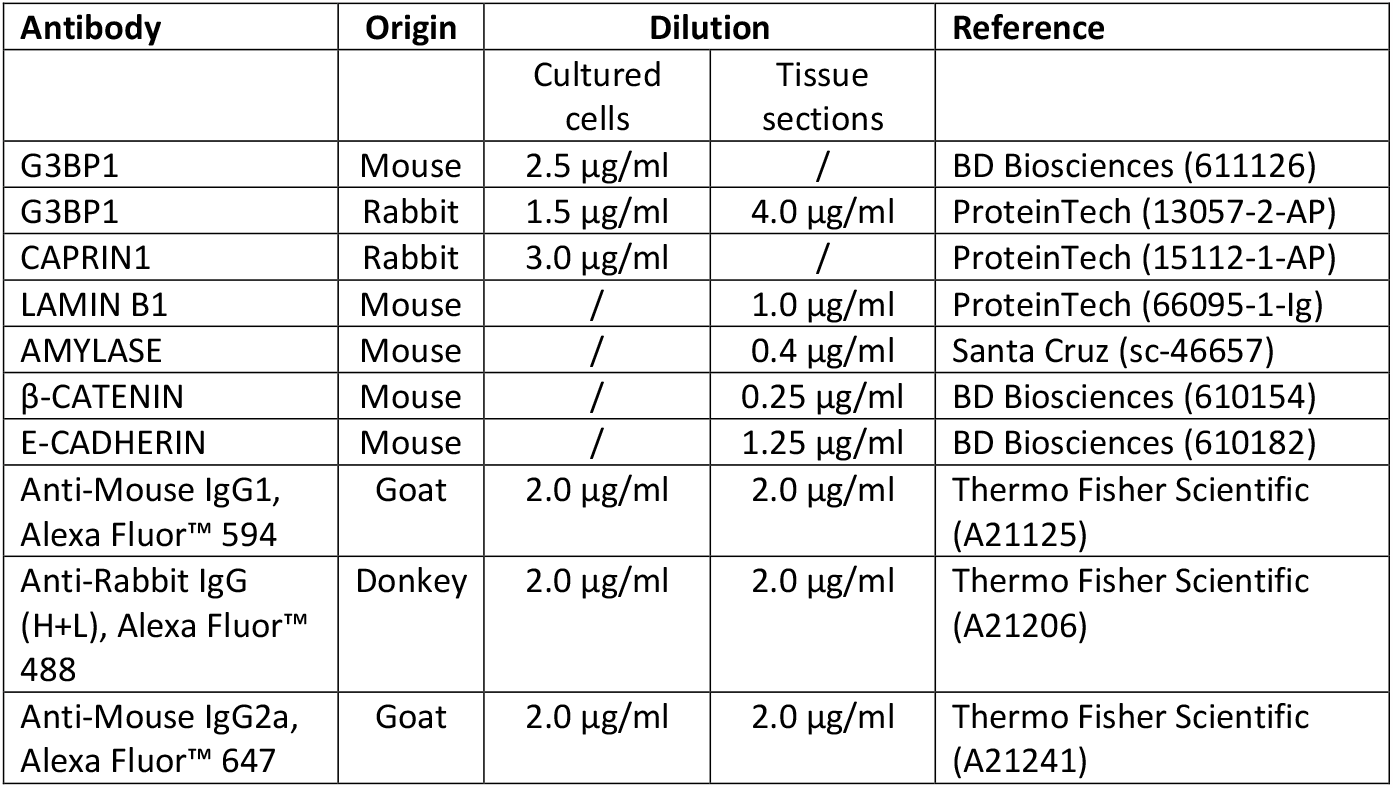

| Antibody | Origin | Dilution |  | Reference |
| --- | --- | --- | --- | --- |
|  |  | Cultured cells | Tissue sections |  |
| G3BP1 | Mouse | 2.5 μg/ml | / | BD Biosciences (611126) |
| G3BP1 | Rabbit | 1.5 μg/ml | 4.0 μg/ml | ProteinTech (13057-2-AP) |
| CAPRIN1 | Rabbit | 3.0 μg/ml | / | ProteinTech (15112-1-AP) |
| LAMIN B1 | Mouse | / | 1.0 μg/ml | ProteinTech (66095-1-Ig) |
| AMYLASE | Mouse | / | 0.4 μg/ml | Santa Cruz (sc-46657) |
| β-CATENIN | Mouse | / | 0.25 μg/ml | BD Biosciences (610154) |
| E-CADHERIN | Mouse | / | 1.25 μg/ml | BD Biosciences (610182) |
| Anti-Mouse IgG1, Alexa Fluor™ 594 | Goat | 2.0 μg/ml | 2.0 μg/ml | Thermo Fisher Scientific (A21125) |
| Anti-Rabbit IgG (H+L), Alexa Fluor™ 488 | Donkey | 2.0 μg/ml | 2.0 μg/ml | Thermo Fisher Scientific (A21206) |
| Anti-Mouse IgG2a, Alexa Fluor™ 647 | Goat | 2.0 μg/ml | 2.0 μg/ml | Thermo Fisher Scientific (A21241) |

### ELISA

Measure of 15-d-PGJ2 levels was conducted using an ELISA kit (ADI-900-023, Enzo Life Sciences) following the manufacturer’s instructions. The measurements were carried out in conditioned medium, 72 hours after seeding 30,000 cells in a 24-well plate.

### Analysis of human patient data

Transcriptomic data normalized in transcripts per kilobase million (TPM) and the Simple Nucleotide Variation datasets for Pancreatic ductal adenocarcinoma (PDAC), Lung adenocarcinoma (LUAD) and Colon adenocarcinoma (COAD) were downloaded from The Cancer Genome Atlas (TCGA) consortium^9^, using TCGAbiolinks v2.14.1 R-package^10^. Only unique primary (01A) tumor samples with available RNA-seq and Simple Nucleotide Variation data were included in the analysis. The tumor samples were categorized on the possible presence of a *KRAS* missense mutation, either in wild-type (*KRAS* WT) or in mutant (*KRAS* MUT). For PDAC: KRAS WT=66; KRAS MUT=99. For LUAD: KRAS WT=364; KRAS MUT=138. For COAD: KRAS WT=241; KRAS MUT=180.

### Statistical analysis

Data were presented as means ± standard error of the mean (SEM). Single comparisons between two experimental groups were done using a paired Student’s t test. To identify significant differences between multiple groups, statistical analyses were performed using a one-way ANOVA followed by Tukey test. For all statistical analyses, the level of significance was set at p < 0.05. Statistical analysis was performed with GraphPad 8.0.2 Prism Software (GraphPad Software Inc., San Diego, CA, United States). *,p < 0.05; **,p < 0.01; ***,p < 0.001; ns, not significant.

## Acknowledgments

We would like to thank members of the laboratory of Patrick Jacquemin and Frederic Lemaigre for help and discussions, Malak Haidar for KPE cells, Jean-Nicolas Lodewyckx and Hajar Dahou for mouse genotyping, and Mourad El Kaddouri and Bruno Maricq for their help in mouse care.

